# Effects of shoe heel height on ankle dynamics in running

**DOI:** 10.1101/2023.08.08.552471

**Authors:** Ali Yawar, Daniel E. Lieberman

## Abstract

Shoes alter the evolved biomechanics of the foot, potentially affecting running kinematics and kinetics that can in turn influence injury and performance. An important feature of conventional running shoes is heel height, whose effects on foot and ankle biomechanics remain understudied. Here, we investigate the effects of 6 – 26 mm increases in heel height on ankle dynamics in 8 rearfoot strike runners who ran barefoot and in minimal shoes with added heels. We predicted higher heels would lead to greater frontal plane ankle torques due to the increased vertical moment arm of the mediolateral ground reaction force. Surprisingly, the torque increased in minimal shoes with no heel elevation, but then decreased with further increases in heel height due to changes in foot posture that are probably a strategy to compensate for potentially injurious ankle torques. We also found that increasing heel heights caused a large increase in the ankle plantarflexion velocity at heel strike, which we explain using a passive collision model. Our results highlight how running in minimal shoes may be significantly different from barefoot running due to complex interactions between proprioception and biomechanics that also permit runners to compensate for modifications to shoe design, more in the frontal than sagittal planes.

## Introduction

During running, the ankle joint transmits large forces and moments, producing the greatest fraction of joint power for propulsion ^1^. Although the ankle primarily rotates in the sagittal plane, its frontal plane kinematics and kinetics play equally important roles in governing ankle stability and preventing injury, most notably sprains from excess inversion or sometimes eversion. The ankle’s behavior during running has been shown to be influenced by the geometric and material properties of shoes, particularly the heel region, which can alter the nature of forces transmitted through the foot and ankle compared to barefoot running ^2,3^. While minimal shoes have sole thicknesses of just a few millimeters with no heel-toe offset, some cushioned shoes have heel heights in excess of 15 mm and similar heel-toe offsets ^4^. Here we investigate how this wide variation in heel heights in running shoes affect ankle stability and motion in the frontal and sagittal planes.

A key destabilizing load in the frontal plane is the torque about the ankle joint, which has contributions from the vertical and mediolateral ground reaction forces (*F*_*z*_ and *F*_*x*_ respectively in figure 1). These forces produce moments proportional to the mediolateral and vertical moment arms about the ankle joint (*r*_*x*_ and *r*_*z*_ respectively in figure 1), both of which are influenced by the size and shape of the shoe heel (figure 1b). In particular, during heel strike in single limb support when the center of pressure (COP) must lie inside the heel base, heel height increases the vertical moment arm, and the width of the heel constrains the maximum mediolateral moment arm. Thus, the height and width of the heel, which tend to be large in modern running shoes, may significantly influence the torque about the ankle joint. Since shoe design features significantly interact with each other, it is difficult to study the effect of heel geometry in isolation, without influence from characteristics like heel cushioning ^4^. Thus, few studies have considered the specific effects of variations in heel geometry and heel toe offset of running shoes on frontal plane ankle dynamics ^3,5^.

**Figure 1.**
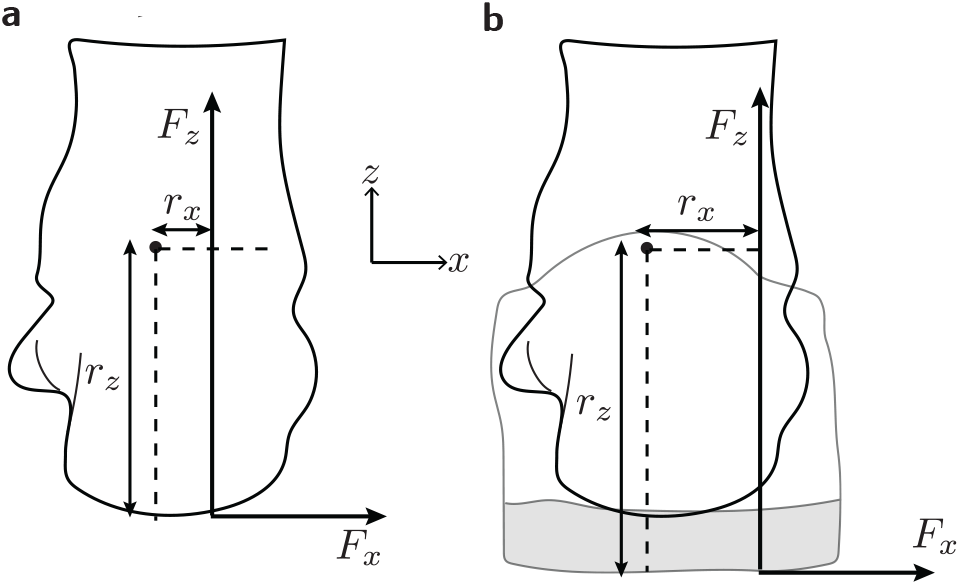
Frontal plane schematic of the foot. **a**, In the frontal plane, the moment arm of the vertical ground reaction force *F*_*z*_ is *r*_*x*_, and that of the mediolateral ground reaction force *F*_*x*_ is *r*_*z*_. **b**, The presence of a shoe increases *r*_*z*_ by the height of the shoe, and increases the possible range of motion of the center of pressure in the mediolateral direction, allowing for potentially larger *r*_*x*_. Figure adapted from Yawar and Lieberman ^2^.

Heel height may also affect the sagittal plane torques and range of motion of the ankle through stance ^3,6,7^. It is unclear if these effects are direct modifications of ankle dynamics or secondary outcomes of changes in stride length ^8^. It is therefore useful to ask how heel height may influence dynamics of the ankle just after landing, specially in rearfoot strike (RFS) runners where the heel is the only point of contact in early stance. When the foot collides with the ground during running, the pre-collision kinetic energy of the lower limb is converted into rotational kinetic energy of the foot to varying degrees depending on strike type ^9^. The vertical ground reaction force acting on the calcaneus in RFS runners may plantarflex the ankle at landing. Further, runners land with forward foot speeds of around 2 m/s ^10^, and so the fore-aft ground collision in rearfoot strike may also contribute to ankle plantarflexion at landing, especially in shoes with elevated heels. Unlike the controlled dorsiflexion of the ankle during forefoot strike landing ^9^, this plantarflexion is likely uncontrolled, as the primary dorsiflexor of the foot, tibialis anterior, does not contract significantly at heel strike in RFS running ^11^. Thus, we predict that heel height influences the early stance plantarflexion velocity of the ankle.

This study measures the effects of small increases in heel heights (up to 26 mm) on the early stance dynamics of the ankle joint in the sagittal and frontal planes during RFS running. We focus our analysis on the first half of stance when we expect heel height to have the most significant effect. We test two hypotheses. The first hypothesis is that an increase in heel height will increase frontal plane torque at the ankle in early stance, due to an increase in the vertical moment arm of the ground reaction force about the ankle. Second, by extending the collision model in Lieberman et al. ^9^ to include a massless heel, we predict that ankle plantarflexion velocity (“foot-slap”) increases with heel height.

## Results

### Frontal plane

There was a significant effect of heel height condition on the mean frontal plane ankle torque (*F*_3,21_ = 29.54, *p* < 0.001, figure 2a). Post-hoc tests showed major differences of 11–32% among the barefoot condition and different heel heights: a 25.3% decrease from barefoot to high heels (*p* < 0.001); 13.2% decrease from low to medium heels (*p* < 0.001); 32.4% decrease from low to high heels (*p* < 0.001); and a 22.1% decrease from medium to high heels (*p* < 0.001). There was a 10.6% increase in the torque from barefoot to low heel (*p* = 0.008).

**Figure 2.**
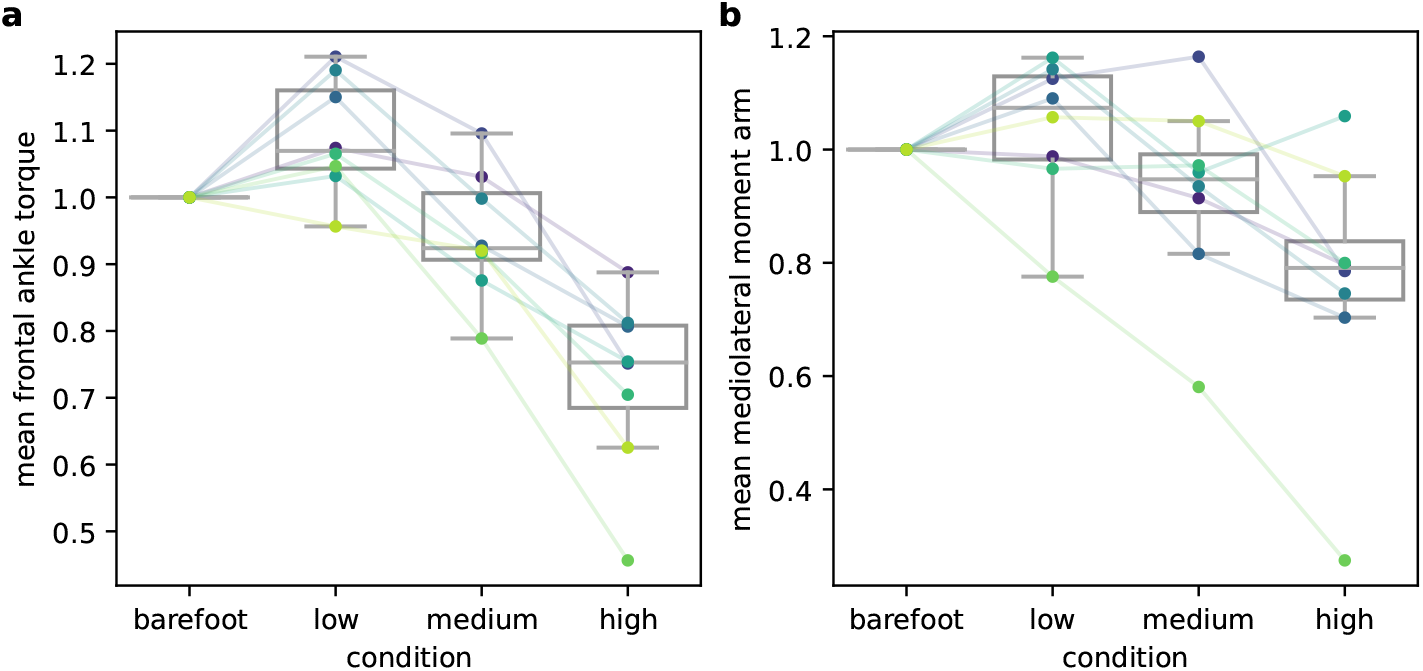
Frontal plane effects of heel height on the ankle. **a**, Box plot of the mean frontal plane ankle torque, shown here normalized by the barefoot condition to highlight relative changes. **b**, Box plot of the mean mediolateral moment arm of the ground reaction force about the ankle normalized by the barefoot condition. Each color denotes one subject’s data.

There was a significant effect of heel height condition on the mean mediolateral moment arm of the ground reaction force about the ankle (*F*_3,21_ = 10.84, *p* < 0.001, figure 2b). Post-hoc tests showed major differences of 11–27% among the barefoot condition and different heel heights: a 22.8% decrease from barefoot to high heels (*p* < 0.001); 11% decrease from low to medium heels (*p* = 0.04); 26.5% decrease from low to high heels (*p* < 0.001); 17.4% decrease from medium to high heels (*p* = 0.004)

### Sagittal plane

From the collision model, the post-collision ankle plantarflexion velocity is 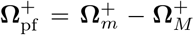 where,

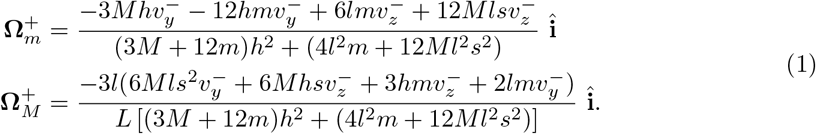

Magnitudes of the barefoot-normalized angular velocity for the foot and shank are plotted in figure 3a, and the barefoot-normalized ankle plantarflexion velocity in figure 3b.

**Figure 3.**
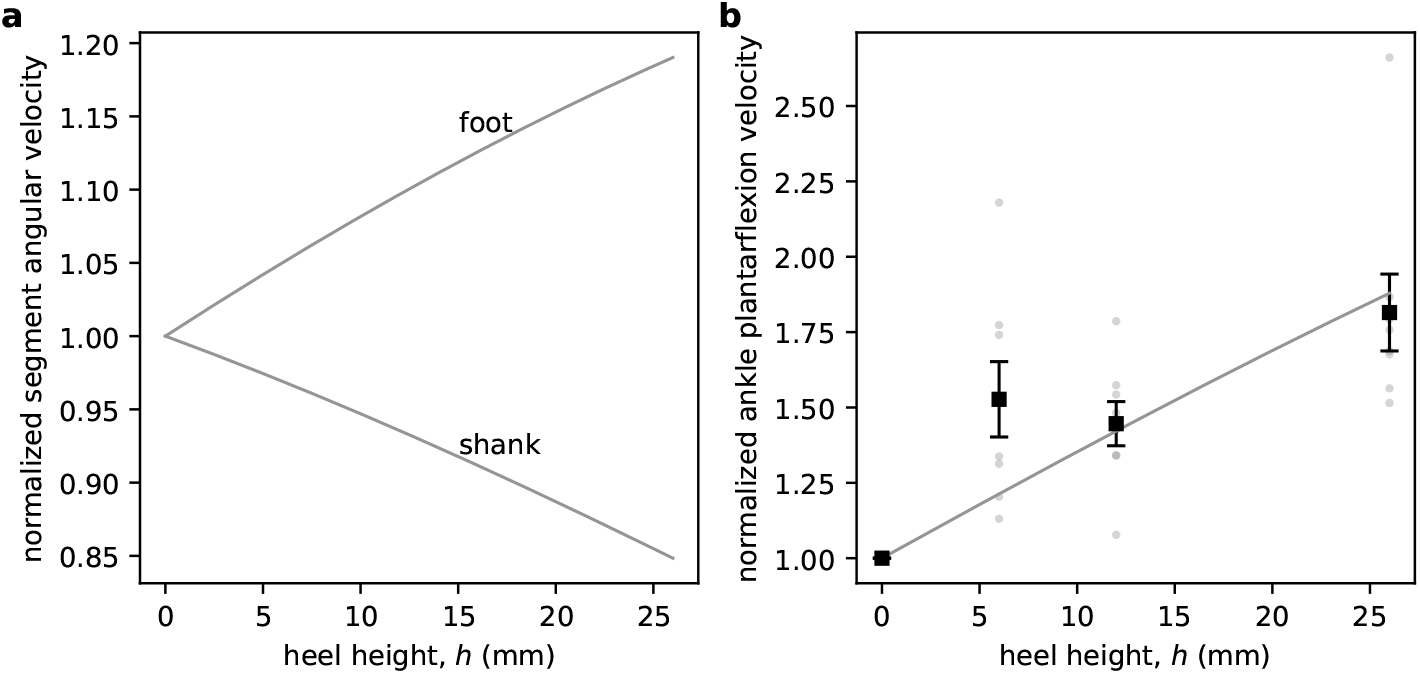
Collision model predictions for ankle plantarflexion velocity. **a**, Magnitudes of foot and shank angular velocities, normalized by the barefoot condition, plotted against heel height *h*. **b**, Scatter plot of normalized ankle plantarflexion velocity against heel heights in different conditions (barefoot: 0 mm; low: 6 mm; medium: 12 mm; high: 26 mm). Black squares show means for each heel height, whiskers show standard error of the mean, and gray dots show individual subjects’ data. The solid gray curve shows the model predicted magnitude of the ankle plantarflexion velocity (foot relative to shank) normalized by the barefoot condition.

There was a large and significant effect of heel height condition on the mean peak ankle plantarflexion velocity just after heel strike (*F*_3,21_ = 49.76, *p* < 0.001, figure 3b). Post-hoc tests showed major differences of 19–75% among all conditions: a 47.5%, 41.8% and 75.3% increase from barefoot to low, medium and high heel conditions respectively (*p* < 0.001 for all comparisons); 18.8% increase from low to high heels, (*p* < 0.001); 23.6% increase from medium to high heels (*p* < 0.001).

## Discussion

Over the last 50 years, there has been a general trend towards higher heels in many but not all running shoes, and today’s runners can now choose between heels that vary from just one or two millimeters to about 40 mm ^23^. Here we tested how these variations affect the kinematics and kinetics of the ankle in the frontal and sagittal planes during running. In order to minimize the influence of shoe features such as bending stiffness and cushioning ^2^, we used stiff heels mounted on minimal running shoes that offered little to no support. Therefore, this study tested effects of just variations in heel height.

As predicted, the magnitude of the frontal plane torque changed significantly with increasing heel heights, but not in accordance with a simple model based solely on the effects of changing vertical moment arms. Notably the torque in the frontal plane was higher in the minimal shoe condition with no heel relative to the barefoot condition, but with increasing heel heights, torque declined (figure 2a). Overall, frontal plane torque in the highest heel condition (26 mm) was over 25% lower than in the barefoot condition. This reduction in torque was a result of reduction in the mediolateral moment arm (*r*_*x*_) of the vertical ground reaction force (*F*_*z*_) about the ankle (figure 2b), as runners tended to run with a center of pressure (COP) closer to the ankle joint in the mediolateral direction over the first half of stance.

While this study was not designed to identify specific mechanisms underlying ankle control, we propose that modulation of foot posture in the frontal plane was the strongest influence on *r*_*x*_. Foot posture has distinct but related effects on the mediolateral positions of the ankle and COP. Across all participants and conditions, the COP was located lateral to the ankle in the entire first half of stance (13.2 ± 4.4 mm, mean ± s.d.). So, in order to reduce *r*_*x*_, either the ankle has to be displaced laterally relative to the COP, or the COP displaced medially relative to the ankle. Once contact is made with the ground, the mediolateral position of the ankle joint is kinematically coupled to the posture of the foot in the frontal plane ^12^. Consistent with this coupling, we observed high step-to-step correlation in the mediolateral displacement of the ankle joint and the eversion angle of the foot within each trial: the more everted (inverted) the foot, the more medially (laterally) displaced the ankle (*r* = 0.93 ± 0.05, mean± s.d across every stance period, figure 4a). Foot posture also affects the location of the COP by influencing the area of contact with the ground within which the COP must lie during the single support phase, although the COP also depends on the position and acceleration of the center of mass. The more inverted the foot, the more lateral the COP is likely to be under the foot ^13^, until it reaches the foot’s lateral boundary. The net effect of these lateral movements of the ankle and COP is that across all conditions and subjects, mean *r*_*x*_ is correlated positively with mean foot eversion (*r* = 0.6, *p* < 0.001, figure 4b). With increasing heel heights, foot eversion angle significantly changed (*F*_3,21_ = 11.55, *p* < 0.001, figure 4c), in a trend similar to that observed in the moment arm and ankle torque (figure 2).

**Figure 4.**
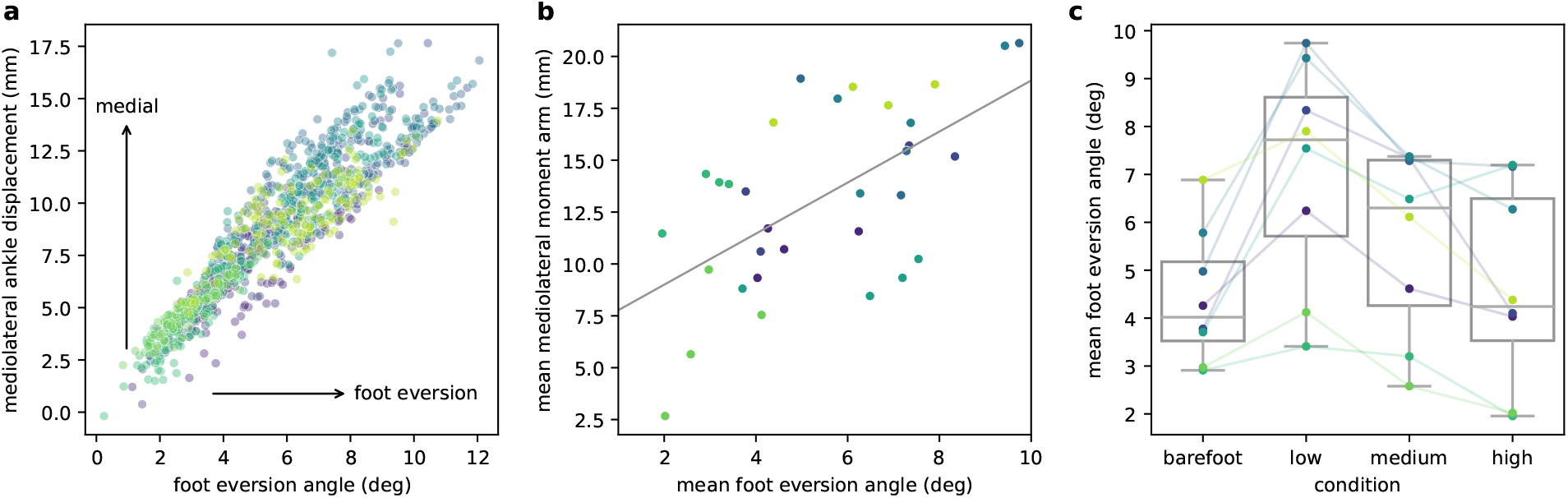
Frontal plane posture of the foot. **a**, Mediolateral ankle displacement from its position at heel strike plotted against the foot eversion angle for every stance period across all subjects. Each dot corresponds to one stance period. **b**, Mean mediolateral moment arm plotted against mean foot eversion angle in each condition and subject. The linear regression line is shown in gray. **c**, Box plot of the mean foot eversion angle over the first half of stance normalized by the barefoot condition. Dots show individual subjects’ data. (Each color denotes the same subject in all three panels).

The ways in which runners adjust foot posture in heels of different heights suggest a possible compensation strategy by RFS runners to protect the ankle from large and potentially injurious frontal plane torques. We did not investigate the specific proprioceptive feedback mechanisms that underlie these adaptations to heel height, but these mechanisms, which presumably include both cutaneous and muscle spindle receptors, deserve further study. It is especially notable that compared to the barefoot condition, even minimal shoes with soles just 6 mm thick led to altered ankle torques in the mediolateral direction. Interestingly, minimal shoes increased ankle torque relative to the barefoot condition, but as heels got higher this torque decreased. While a minimal shoe likely reduces feedback from cutaneous sensors ^14^, we hypothesize that the additional vertical moment arm associated with higher heels could perhaps more effectively engage muscle spindles or golgi tendon organs in the muscles of the leg that control inversion and eversion, notably the posterior tibialis and peroneus longus. Thus, while it is generally accepted that minimal shoes affect foot kinetics and kinematics less than conventional shoes ^2,15^, our results highlight that there are still salient differences between running barefoot and in minimal shoes whose consequences may not be completely understood and deserve further study.

Higher heels also have significant effects in the sagittal plane. In particular, using heel heights within the range of those typically found in running shoes, we found that higher heels were associated with faster ankle plantarflexion velocities in early stance, with the highest heel condition showing an almost 75% increase in peak plantarflexion velocity compared to barefoot. This effect is likely passive, and it is predicted reasonably well by a simple model of collision at heel strike extended from Lieberman et al. ^9^ (figure 3). The pre-collision translational kinetic energy of the foot and shank is partially converted into rotational kinetic energy of the two segments post-collision as they rotate forwards in the sagittal plane. With increasing heel heights, shank angular velocity decreases and foot angular velocity increases (figure 3a), resulting in a net increase in ankle plantarflexion velocity (figure 3b). The biomechanical effects of high ankle plantarflexion velocities are not well studied but repetitive, fast plantarflexion of the foot during running may contribute to overuse injuries of the tibialis anterior muscle and tendon, including chronic exertional compartment syndrome in the lower leg ^16,17^. Future studies are necessary to test this hypothesis, but we note that chronic exertional compartment syndrome of the anterior tibialis has been shown to be prevented or treated by switching from a rearfoot strike in conventional shoes which presumably involve rapid plantarflexion right after heel strike to a forefoot strike which eliminates this motion ^18^.

Our findings have implications for future studies on shoe design and how runners adapt to higher heels. Currently, data do not exist to evaluate the extent to which runners adapt differently to higher heels in terms of controlling the center of pressure or the rate of plantarflexion, or if there is a critical heel height above which foot posture cannot mitigate high frontal plane torques. Longer duration studies with minimal shoes would be useful to evaluate if frontal plane ankle torques reduce with continued use or training, shedding more light on the function of cutaneous feedback from the foot sole. In people with tibialis anterior paresis, rapid ankle plantarflexion at heel strike (or “drop foot”) leads to spatiotemporal gait modifications ^19^. It could be useful to evaluate if using minimal shoes or shoes with shorter than usual heels can reduce the initial ankle plantarflexion velocity and a return to pre-paretic gait. Finally, shoes that modulate foot posture, and therefore mediolateral center of pressure, might help runners manage ankle instability injuries.

The effects of variations in heel height may not be limited to the ankle but also extend to the knee and overall gait. For example, a higher heel height (more precisely the heel-toe offset) leads to a more plantarflexed ankle, and gives rise to a tradeoff between loading at the ankle and the knee ^7,20^. Higher heels may also increase stride length during running ^8^, which may in turn influence the rate and magnitude of vertical ground reaction forces at heel strike ^21^. Similar to these effects, the changes in foot posture, frontal plane torque, and plantarflexion velocity that we observe at the ankle are likely to alter the kinetics and kinematics of joints further up the kinematic chain, and may give rise to tradeoffs that may be exploited to improve running economy and manage injuries.

This study has several limitations. We used heels made of a relatively rigid material compared to the materials used in conventional running shoes. The difference between the loaded and unloaded heights of a compliant shoe heel must be taken into account when evaluating its effects on foot biomechanics, specially since for a given material, higher heels are more compliant ^22^. Heel widths in this study were approximately 60 mm, which is about 25 -40% smaller than the width of some conventional shoe heels ^23^. The width of the heel constrains the mediolateral range of motion of the COP. Since wider heels allow a greater mediolateral range of motion of the COP, runners’ strategies to control ankle torque might differ from what we observed. Finally, subjects did not run at self selected speeds, and since each trial duration was 30 s, any gait adaptations or effects of fatigue were not present.

## Methods

### Participants

We recruited *N* = 8 participants (6 males, 2 females; age 29.5± 7.1 years, body mass 82.9± 14.4 kg, leg length 0.92 ± 0.06 m, mean± s.d.) with no recent history of lower limb injury. Participants provided written informed consent, and the study was approved by the Institutional Review Board at Harvard University.

### Experimental protocol

Subjects ran at Fr = 1 on an instrumented treadmill (Bertec, Columbus, OH, USA). The average speed across participants was 3.0± 0.1 m/s (mean ± s.d.). Subjects were asked to heel strike, and strides with non heel-strikes (judged by the absence of a transient peak in the vertical ground reaction force within the first 20% of stance) were removed from the analysis. The trials were conducted in four conditions in a randomized order: barefoot, in zero-drop minimal shoes of 6 mm sole thickness (“low”), minimal shoes with an additional 6 mm heel (“medium”), minimal shoes with an additional 20 mm heel (“high”). All shoes were zero-drop minimal shoes (Merrell Rockford, Michigan, USA, model: Vapor Glove, US Men’s size 10) modified by a professional cobbler (Felix Shoe Repair, Cambridge, MA, USA) who added custom designed heels (for more details see Addison and Lieberman ^22^). The material used to construct heels had a Young’s modulus of 32 MPa.

In the barefoot condition, three markers were placed on the calcaneus following the Leardini marker set ^24^, and a fourth marker on the head of the second metatarsal. In the shod conditions, four markers were placed directly on the shoe approximately aligned with the markers used in the barefoot condition, following the guidelines for shoe mounted markers ^25^. The rearfoot markers were used to define a foot segment. Additional markers were placed on the medial and lateral malleoli, the tibial tuberosity, and the medial and lateral femoral epicondyles. A four marker cluster was also placed on the shank. The mid-point of the two malleoli markers was defined as a virtual ankle joint.

### Data collection and analyses

Kinematic data were collected using an 8-camera motion capture system at 200 Hz (Qualisys AB, Gothenburg, Sweden). Kinetic data were recorded at 2000 Hz. Kinematic and kinetic data were low-pass filtered using fourth and eighth order Butterworth filters respectively, with a common cutoff frequency of 25 Hz. Data were initially processed in Visual3D (C-Motion, Inc., Moyds, MD, USA). Further data processing and analysis was performed in MATLAB 9.11.0.1837725 (Natick, MA, USA) and Python 3.1. A total of 32 trials were analyzed for 8 subjects and 4 conditions each. The laboratory and all joint coordinate axes are defined such that *x* is along the medio-lateral direction, *y* is along the antero-posterior direction, and *z* is along the superior-inferior direction. Each 30 s trial was segmented into intervals from touchdown to toe off (stance) using *F*_*z*_ = 100 N as the threshold for foot contact detection, and the stance time was normalized to be between 0-100%. The mean mediolateral moment arm is defined as the difference between the *x* components of the center of pressure and the ankle joint position in the lab frame averaged over the first 50% of stance. The mean frontal plane ankle torque is defined as the cross product of the frontal plane ground reaction force with the frontal plane moment arm about the ankle averaged over the first 50% of stance. Ankle plantarflexion velocity in the sagittal plane is defined as the maximum *x* component of the ankle joint angular velocity expressed in the shank coordinate system in the first 20% of stance. The rotation of the foot segment is measured with respect to the lab frame following an intrinsic *x*–*y*–*z* Euler angle scheme, and normalized within each stance period to the foot segment at the moment of heel strike. The mean foot eversion angle is taken to be angle of rotation about the *y* axis averaged over the first 50% of stance. Mediolateral ankle displacement is defined as *x* coordinate of the virtual ankle joint with respect to its position at heel strike, and is averaged over the first 50% of stance. Measurements are further averaged over all stance periods in a trial before performing statistical analyses.

### Collision model

#### Notation

We follow the notation conventions used in Lieberman et al. ^9^. Scalars are denoted by lowercase italics letters, and vectors by bold letters, with subscripts denoting the body or landmark associated with each vector. Unit vectors along the *x, y* and *z* axes are denoted by 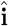, 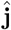 and 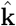 respectively. Moment vectors have subscripts “/A” to denote the point A about with the moment is defined. Quantities just before the collision are labeled with superscript ‘-’ and those just after with superscript ‘+’. Subscripts *m* and *M* refer to the foot and shank segments respectively, and *m* + *M* refers to the two segments together. This analysis is restricted to the sagittal plane (*yz*), and the direction of running is along the positive *y* axis.

We define the plantarflexion velocity of the ankle as the angular velocity of the foot with respect to the shank, just after collision with the ground. Figure 5 shows a schematic of a sagittal plane model of the lower limb which is adapted from Lieberman et al. ^9^. The shank has length *L*, mass *M*, moment of inertia **I**_*M*_, and center of mass at point E; the foot has length *l*, mass *m*, moment of inertia **I**_*m*_ and center of mass at point D. Segment moments of inertia are defined about the respective centers of mass. The foot and shank are connected via a frictionless hinge at point B, which models a zero stiffness ankle. The heel is defined as the fraction of the foot’s length posterior to the ankle joint with length *sl* and a massless vertical projection of height *h*.

**Figure 5.**
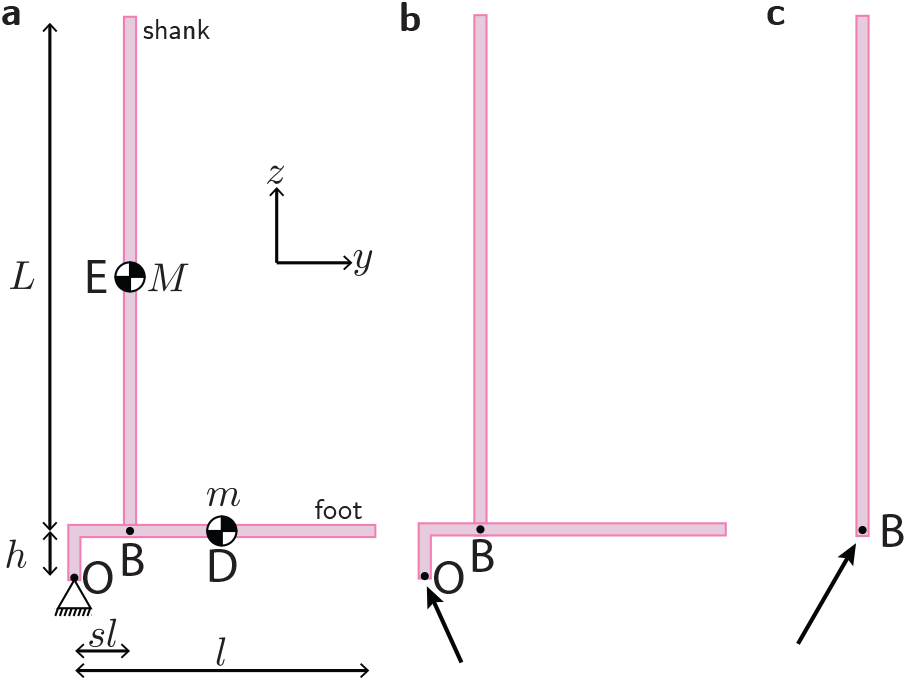
Schematic and free body diagrams of the lower limb model. **a**, The lower limb is modeled with a shank of mass *M* and length *L*, and a foot of mass *m* and length *l* connected at a frictionless hinge at point B which models the ankle joint. The heel extends by a length *sl* posterior to the ankle and has a massless vertical projection of height *h*. The centers of mass of the foot and shank segments are at points D and E respectively, and the ground contact occurs at point O. The direction of running is along the *y* axis in this view. **b** Free body diagram of the entire system showing the reaction force at the ground contact point O. Finite forces like gravity are ignored in the model and not shown here. **c**, Free body diagram of the shank alone, showing the reaction force at the hinge B.

We assume that just before collision, the entire lower limb is moving with the velocity 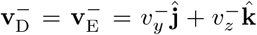 and with zero initial angular velocity 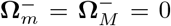. The foot and shank system collides with the ground at point O, and the post collision angular velocities of the foot and shank segments are 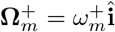 and 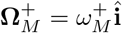. Following Lieberman et al. ^9^, we assume a rigid, plastic collision at point O, so that the collision is instantaneous, the configuration of the system does not change during the collision, and the point O comes to rest after the collision. At any time before, during or after collision, the angular momenta of the shank about the point B and the whole system about the point O are given by,

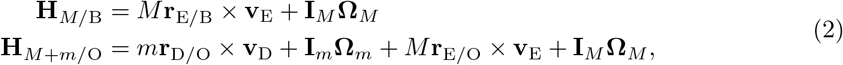

where

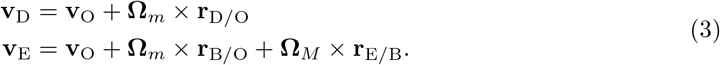

Since the collision is assumed to be instantaneous, the contact forces acting at points B and O are infinitely large. Finite forces such as gravity are negligible compared to the contact forces, and they contribute no torque-impulses in the collision. Thus, there are no torque-impulses about the points B and O, and we can write angular momentum balance for the shank segment about point B, and for the whole system about point O,

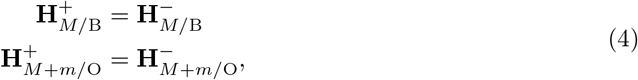

where only the *x*-components the angular momenta are non-zero, resulting in two independent equations. Using these equations we solve for the two unknowns, 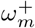 and 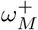 which are the post-collision foot and shank angular speeds respectively, and thus obtain the post collision ankle plantarflexion velocity 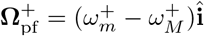.

#### Model parameters

Shank length *L* is assumed to be half the mean leg length of the participants. We use *l* = *L/*1.53, *M* = 4.5% and *m* = 1.4% of the mean participant body weight^26^. The heel to foot length ratio *s* = 0.21 based on published radiographic data^27^. The pre-collision foot velocity components were assumed to be *v*_*y*_ = 1.38 m/s and *v*_*z*_ = *−* 0.82 m/s using the mean fore-aft and vertical velocity of the ankle joint at heel strike across all subjects and conditions (since the foot and shank are assumed to move with the same velocity just prior to collision).

#### Statistical methods

We performed a Type-III one-way analysis of variance (ANOVA) with Satterthwaite’s method to test the effect of heel condition on various measured quantities. First, we created linear mixed models with heel condition as the fixed factor (four levels: barefoot, low heel, medium heel, high heel), and subject as random factor, given by

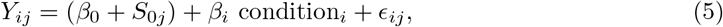

where *i* = 1 舰 3 is the heel condition index and *j* = 1 舰 8 is the subject index. Thus, the dependent variable *Y* has 32 values. The model estimates are the global intercept *β*_0_ corresponding to the barefoot condition (*i* = 0), and the fixed effects coefficients *β*_*i*_ corresponding to the three shod conditions (*i* = 1 舰 3). The random intercepts *S*_0*j*_ quantify inter-subject variability and are assumed to be normally distributed with zero mean. The overall model residuals *E*_*ij*_ are also assumed to be normally distributed with zero mean. If the ANOVA was significant (at *α* = 0.05), we performed post-hoc pairwise comparisons with Benjamini-Hochberg adjustments to the *p*-values. Statistical tests were performed in R 4.1.2 ^28^, using the `lme4` ^29^ and `multcomp` packages ^30^.

## Acknowledgements

We thank Brian J. Addison, and Christos Soillis (Felix Shoe Repair, Cambridge, MA) for designing the modified shoes. A.Y. thanks the American School of Prehistoric Research for funding support.

## Author contributions statement

D.E.L. and A.Y. conceived the experiment; A.Y. conducted the experiment, analyzed the data and performed the collision calculation in consultation with D.E.L.; A.Y. and D.E.L. interpreted the results and wrote the manuscript.

